# Flipper: An advanced framework for identifying differential RNA binding behavior with eCLIP data

**DOI:** 10.64898/2026.03.13.711628

**Authors:** Keegan Flanagan, Shuhao Xu, Gene W. Yeo

## Abstract

**Motivation:** Crosslinking and immunoprecipitation (CLIP) methods are widely used to identify RNA binding protein (RBP) binding sites and assess how treatments alter RBP binding patterns and regulatory activity. However, current tools for differential RBP binding analysis lack core features required for rigorous statistical inference, including proper normalization and appropriate handling of replicate experiments. Furthermore, existing approaches cannot adequately separate expression or splicing driven effects from true changes in RBP binding.

**Results:** Here we present Flipper, an application purpose-built for the analysis of differential RBP binding. Flipper introduces several innovations that adapt the DESeq2 framework for eCLIP data. These include the use of input controls to account for expression and splicing driven effects and hierarchical normalization that adjusts for technical variation without confounding signal to noise ratios. We demonstrate that Flipper exhibits high specificity when applied to real differential eCLIP data while also providing deeper biological insights. In addition, analyses of both real and simulated data indicate that Flipper outperforms existing approaches. Together, these results highlight Flipper as an effective framework for differential RBP binding analysis.

**Availability:** Flipper and its tutorials are available at https://github.com/YeoLab/flipper and https://doi.org/10.5281/zenodo.21136504.

## 1 Introduction

RNA binding proteins (RBPs) modulate a wide range of cellular processes (Gerstberger et al., 2014; Hentze et al., 2018), including translation (Svitkin et al., 1996), RNA degradation (Hasan et al., 2014), and splicing (Lovci et al., 2013). As a result, mutations or other defects in these proteins have been implicated in numerous genetic diseases and cancers (Verkerk et al., 1991; Bardoni et al., 2001; Lukong et al., 2008; Kechavarzi and Janga, 2014). Consequently, studies seeking to perturb RBP function through mutations or drug treatments have become increasingly common (Li and Kang, 2023; Bertoldo et al., 2023), creating an urgent need for experimental and computational frameworks for studying changes in RBP binding.

Investigations of RBP–RNA interactions have long relied on cross-linking and immunoprecipitation (CLIP) methods. Although there are several variations of CLIP (Hafner et al., 2021), they share a common experimental framework. RBPs are cross-linked to their RNA targets, and antibodies are used to immunoprecipitate the RBP of interest along with its associated RNA. The recovered RNA is then sequenced, and genomic regions with enriched read coverage are identified as putative binding sites using peak calling algorithms such as Piranha (Uren et al., 2012), DEWSeq (Schwarzl et al., 2024), and Skipper (Boyle et al., 2023).

Early approaches for differential analysis using CLIP relied on performing peak calling in two conditions and designating peaks unique to each condition as differential. While straightforward, the qualitative nature of this analysis has major shortcomings. This approach is hereafter referred to as the ad-hoc method. Quantitative alternatives include repurposing peak calling software for differential analysis (Uren et al., 2012) as well as published differential CLIP tools such as dCLIP (Wang et al., 2014) and DeepRNA-reg (Sekhon et al., 2025). Although these methods represent conceptual improvements over the ad-hoc approach, they also have limitations. For example, dCLIP and DeepRNA-reg are unable to accommodate multiple CLIP replicates, while repurposed peak callers were not originally designed for differential inference. Furthermore, all these methods lack a key feature required for rigorous differential binding analysis: control for RNA substrate availability.

The number of RNA fragments recovered by CLIP at a given genomic locus reflects both the strength of RBP–RNA binding and the availability of the RNA substrate. This availability depends on the abundance of the transcript and whether the locus is retained or removed from the RNA by splicing. Many perturbations that influence RBP binding also alter substrate availability through changes in expression or alternative splicing. Without a strategy to disentangle these effects, changes in pulldown cannot be unambiguously attributed to altered binding rather than altered substrate availability.

Most CLIP protocols do not provide direct information on RNA substrate availability, requiring additional RNA-seq data to control for availability. Even then, careful consideration is required to ensure that RNA-seq and CLIP measurements are comparable. Enhanced CLIP (eCLIP) (Van Nostrand et al., 2016) offers a solution, as it includes a size-matched input (IN) fraction alongside the immunoprecipitate (IP) fraction. Although this IN fraction is traditionally used for background correction, it can also be leveraged to control for RNA substrate availability. In this study, we present Flipper, a Snakemake pipeline that integrates IP and IN count data within a unified framework, enabling explicit modeling of changes in RBP binding while accounting for underlying RNA substrate availability. Specifically, Flipper extends the DESeq2 (Love et al., 2014) methodology to test interactions between IP and IN counts across conditions, allowing differential binding to be interpreted as changes in pulldown normalized to RNA substrate availability rather than changes in IP signal alone. DESeq2 has been previously applied to the analysis of eCLIP data (Schwarzl et al., 2024) and to differential analysis of similar assays such as methylated RNA immunoprecipitation sequencing (MeRIP-seq) (Tang et al., 2021). However, DESeq2 has yet to be used for differential eCLIP analysis.

Flipper adapts DESeq2 for differential eCLIP analysis by introducing several innovations. These include aggregating IN counts by genomic feature for increased statistical power and implementing a hierarchical normalization strategy that independently accounts for pulldown efficiency and sequencing depth in the IP fraction. For computational efficiency, Flipper restricts analysis to genomic windows identified by the Skipper peak caller (Boyle et al., 2023) as significantly enriched in at least one condition. Together, Skipper and Flipper form a complete end to end pipeline for differential eCLIP analysis. Using both real and simulated datasets, we demonstrate that Flipper recapitulates established differential RBP binding results while consistently outperforming existing approaches.

## 2 Materials and methods

### 2.1 Data preprocessing and window-level quantification

Flipper operates directly on the outputs of the Skipper peak caller (v2.0). Skipper was run using the reference genome and gene annotation from GENCODE v49. Before window construction, the annotation was filtered using the workflow provided with Skipper v2.0, which prioritizes transcripts based on GENCODE transcript-quality tags. Skipper then partitions the transcriptome into approximately 100 base pair windows while ensuring that windows do not cross feature boundaries (e.g. intron-exon junctions). For each window, Skipper quantifies IP and IN reads and applies a beta-binomial framework to identify windows in which IP read counts are significantly enriched over IN. For additional details on Skipper, see the original publication (Boyle et al., 2023) and documentation at https://github.com/yeolab/skipper.

Flipper then constructs a unified dataset containing IP and IN read counts from windows with significant IP enrichment in at least one treatment group (Fig. 1A). We initially attempted to use window-level IN values to control for RNA substrate availability, as in MeRIP-seq differential analysis (Tang et al., 2021; Guo et al., 2022). However, MeRIP-seq uses total RNA as the IN (Meyer et al., 2012), while eCLIP uses size-matched IN that has undergone the same size selection as the IP (Van Nostrand et al., 2016). As a result, eCLIP IN is sparser than MeRIP-seq IN, making window-level IN counts too low to reliably estimate changes in RNA substrate levels.

**Figure 1:**
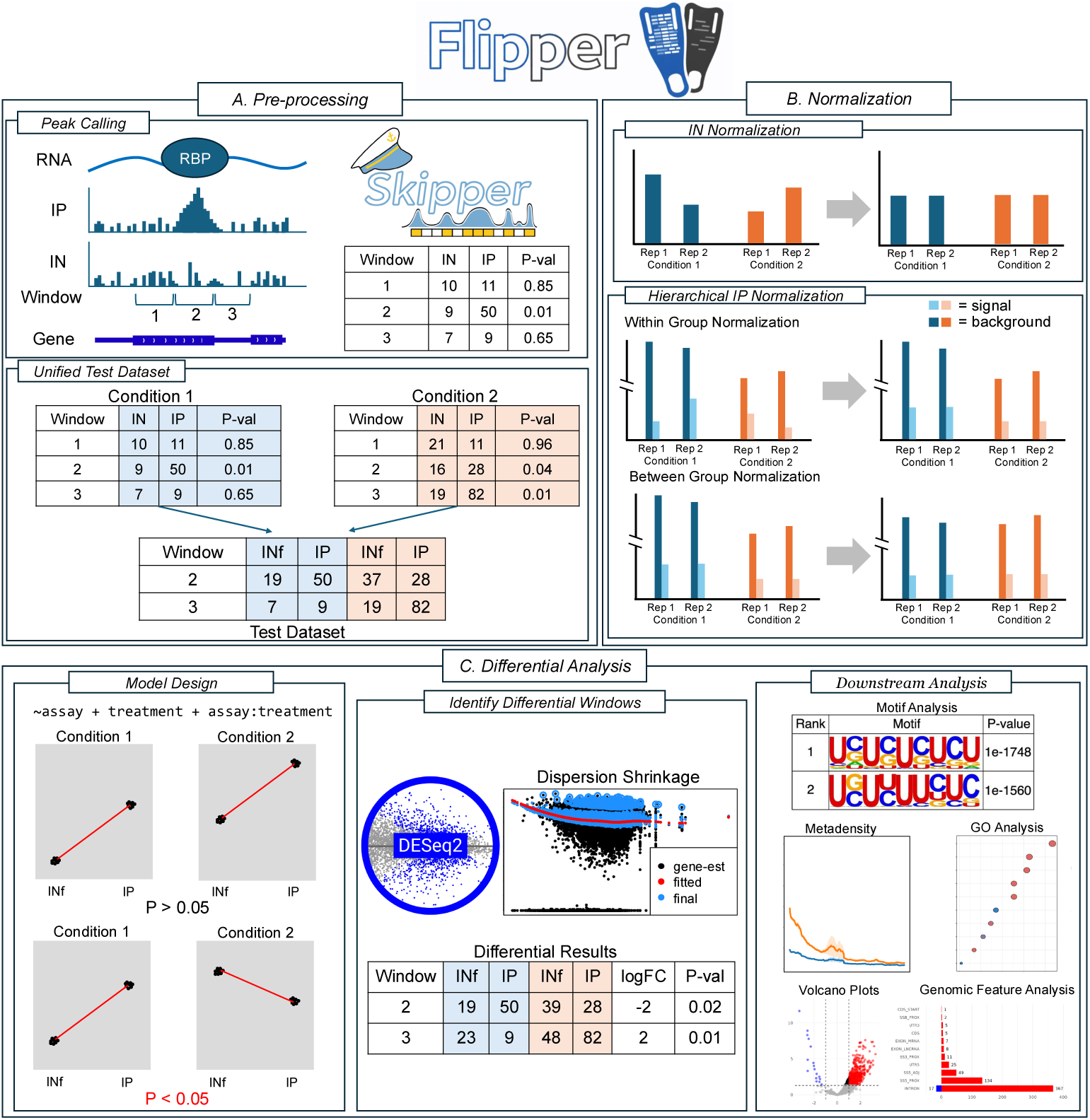
Overview of Flipper. The workflow consists of three major stages. A) Preprocessing and peak consolidation. Candidate binding sites are identified in each condition using Skipper and merged into a unified count matrix containing window-level IP counts and feature-level input (INf) counts. B) Independent normalization of INf and IP signals. INf counts are normalized together in one step, whereas IP counts undergo hierarchical normalization to separately account for enriched and background regions. C) Differential analysis using a DESeq2 framework that models assay type and condition to detect changes in IP signal relative to INf between treatments. Significant sites are subsequently summarized through downstream analyses. Alt text: Diagram depicting the main steps of the Flipper workflow. The workflow is split into three subpanels labeled a–c, covering preprocessing with Skipper, hierarchical normalization, and statistical analysis with DESeq2.

Therefore, Flipper aggregates IN read counts across all windows within each genomic feature (e.g. intron, exon, or UTR) or gene to obtain a feature-level (INf) or gene-level (INg) input value. These aggregate IN values are then used for substrate availability control in downstream analyses. INf is the default mode of Flipper, as it generally produces more conservative and accurate results than INg. The benefits and tradeoffs of each mode are discussed in greater detail in section 3.3 of the main text.

### 2.2 Normalization framework

The assumption of normalization methods applied to traditional differential RNA-seq analysis is that most genes will remain unaffected, allowing identification of a discrete subset of treatment affected genes (Love et al., 2014). This assumption often fails in differential CLIP analysis because a mutation in an RBP binding region or the attachment of a small molecule can cause near global changes in binding affinity and pulldown efficiency. One possible solution is to use non-binding background regions to control for sequencing depth (Stark et al., 2011); however, this strategy does not account for differences in signal to noise ratios between IP replicates (Shao et al., 2012). Technical variation can affect both background and true binding signal, which can cause discordance in the signal to noise ratios across samples and replicates.

For example, variation in washing efficiency may increase background read counts in one IP replicate relative to another, whereas variation in pulldown efficiency may reduce reads at true binding sites in that same replicate. Normalizing only to background reads can therefore exaggerate apparent binding differences, increase variability, and reduce sensitivity (Fig. S1). To address this problem, we developed a two stage hierarchical normalization strategy (Fig. 1B). IP data are partitioned into background regions and binding regions identified by Skipper. In the first normalization step, scaling factors or offsets are estimated from binding regions within individual treatment groups using either median of ratios normalization (MOR) (Love et al., 2014) or EDAseq (Risso, 2011). This step assumes that overall binding levels are comparable among replicates within each treatment group. Because these replicates represent the same biological condition and are processed using the same experimental protocol, systematic differences in global binding are not expected. In the second normalization step, scaling factors are estimated from background regions across all treatment groups and replicates using MOR. These background derived scaling factors are then averaged within each treatment group and combined with the binding derived scaling factors. This procedure accounts for differences in sequencing depth between treatment groups without imposing assumptions about signal to noise ratios.

In contrast to IP libraries, IN libraries do not undergo immunoprecipitation and therefore do not exhibit binding-dependent signal to noise variation. As such, all IN samples are normalized together using a conventional application of MOR normalization (Love et al., 2014).

Although the EDAseq method offers additional normalization options, including adjustments for gene length and GC content, its increased complexity makes it difficult to guarantee that all available normalization options are appropriate for every dataset. As such, MOR is used as the default normalization method in Flipper. All results presented in this study were generated using hierarchical MOR normalization.

### 2.3 Differential binding analysis with DESeq2

The unified table containing both IP and feature-level input (INf) values for each window is used as input for DESeq2 (v1.46.0) using the following design formula:

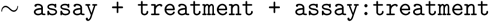

where assay indicates whether counts originated from IP or INf, and treatment indicates whether counts are from the control or treatment group (Fig. 1C). This interaction design tests for changes in the relationship between IP and INf across conditions rather than testing changes in IP alone, thereby controlling for treatment associated changes in RNA substrate captured by INf.

Normalization is implemented using either the sizeFactors or normalizationFactors functions, depending on whether normalization was performed using MOR or EDAseq, respectively.

Windows with an adjusted p-value below 0.05 and an absolute log_2_ fold change greater than 1 were considered significantly differentially bound. The log_2_ fold change corresponds to the assay:treatment interaction term and represents the change in the IP/INf ratio between treatment groups.

### 2.4 Comparison with existing differential binding methods

We benchmarked Flipper against four differential binding strategies: (i) an ad-hoc peak comparison method, (ii) a repurposed peak caller approach, and (iii–iv) two published tools, dCLIP (Wang et al., 2014) and DeepRNA-reg (Sekhon et al., 2025). There exist many variants of the first two approaches. Nearly any peak caller can be used in an ad-hoc framework, and any peak caller that incorporates IN background can, in principle, be repurposed for differential analysis. We used Skipper for both approaches, as Flipper also operates on windows defined by Skipper, thereby maintaining consistent definitions of binding windows across the ad-hoc, repurposed peak caller, and Flipper methods.

For the ad-hoc method, we directly analyzed the same Skipper runs used by Flipper, classifying windows exclusive to the control group as decreased binding and those exclusive to the treatment group as increased binding. For the repurposed Skipper approach, Skipper’s IP and IN inputs were replaced with IP data from the treatment and control groups. Because Skipper performs a one-directional test, each comparison required two runs: treatment IP was tested against control IP to identify increased binding, and control IP was tested against treatment IP to identify decreased binding. This specific repurposing of the Skipper peak caller will be referred to as Diff-Skipper from this point onward.

Both dCLIP v1.0.0 and DeepRNA-reg v1.0 were downloaded from GitHub and run with default parameters. Both programs used the deduplicated BAM files generated by Skipper as input, with dCLIP requiring conversion to SAM format. For both tools, experimental replicates were concatenated into a single alignment file per condition according to the recommendations of the dCLIP and DeepRNA-reg authors. DeepRNA-reg additionally requires a BED file specifying genomic regions to test. We chose to apply DeepRNA-reg only to genomic windows found to be significant binding sites in at least one treatment group by Skipper to maximize its comparability to Flipper.

Output formats and reported statistics differ substantially across methods. Comparisons were therefore limited to metrics available across all tools, such as the total number of differential regions identified. Estimates of differential binding strength or statistical certainty were not compared, as the metrics produced by each program are not directly analogous and are often absent from one or more methods. For example, neither DeepRNA-reg nor dCLIP provides quantities analogous to p-values.

### 2.5 Differential RBP binding datasets

To date, there are no gold standard differential eCLIP datasets in which all true differential binding events are known. As a result, evaluation of differential binding methods must rely on datasets that provide some inherent expectations rather than a complete ground truth. These expectations may arise from prior experimental characterization of a perturbation, a strong mechanistic prediction, or inactive-compound and vehicle controls that provide internal negative comparisons. Based on these criteria, three diverse eCLIP datasets were selected for analysis.

The first dataset was generated from an eCLIP experiment performed on the RBP NONO in 22Rv1 human prostate cancer cells (Kathman et al., 2023b,a). Cells were treated with the small-molecule compound R-SKBG-1, which is reported to associate with NONO and stabilize RBP binding, as well as with a corresponding inactive compound (the enantiomer S-SKBG-1) and a DMSO vehicle control. This design provides both a predicted direction of effect and two negative control comparisons. Two replicates were collected for each of the three conditions. Raw FASTQ files are available through GEO under accession GSE198212.

The second dataset was generated from an eCLIP experiment performed on the RBP DDX42 in HCT116 cells (Lazear et al., 2023a,b). Cells were treated with the small-molecule compound WX-02-23, which was designed to perturb splicing activity and DDX42 binding preferences, as well as with a corresponding inactive compound (WX-02-43) and a DMSO vehicle control. The inactive compound and vehicle conditions provide internal controls for effects specific to WX-02-23. Two replicates were collected for each of the three conditions. Raw FASTQ files are available through GEO under accession GSE220185.

The third dataset was generated from an eCLIP experiment performed on the RBP PUF60 in MDA-MB-231 cells (Tankka et al., 2025a,b). Cells were made to express either PUF60 carrying the L140P mutation in its RRM1 domain or wild type PUF60. The mutation was previously reported to decrease binding to the canonical PUF60 motif (Kralovicova et al., 2020). Two replicates were collected for each condition. Raw FASTQ files are available through GEO under accession GSE280899.

### 2.6 Quantification of intron retention

Each IN fraction BAM file was analyzed separately using IRFinder v2.0.1 (Lorenzi et al., 2021). IRFinder was run using the same reference genome and filtered gene annotation used by Skipper and Flipper. Intron retention was evaluated using IRFinder output metrics, including IRratio. The reported metrics and their definitions are provided in Supplementary Table 3 and its associated caption. All other IRFinder parameters were left at their default values.

### 2.7 Simulation framework

Simulated read counts were generated for 40,000 genes across 8 replicates using negative binomial distributions following the approach described in Love et al. (2014). The number of genes simulated reflects a rough estimate of the total number of unique genes, coding or non-coding, observed in the human genome (Salzberg, 2018). For each gene *i*, a baseline expression coefficient *α_i_* was sampled from a normal distribution, and the mean counts were defined as 2*^αi^*. Gene specific dispersions were computed according to the function of the total read counts. Replicates were divided evenly between control and treatment conditions, with treatment samples having their mean counts modulated by a treatment coefficient *β_i_*. *β_i_* was sampled from a normal distribution for each gene and was subsequently integrated into the expected mean counts 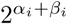. Read counts for each replicate were then generated as independent draws from a negative binomial distribution using the mean and dispersion values for each gene. This allows each gene to have a shared underlying abundance while also allowing replicate to replicate variability and condition specific effects. The resulting gene-level counts were subsequently downsampled to more closely reflect the sparsity observed in eCLIP data.

Gene lengths for the simulation were drawn directly from the distribution of gene lengths seen in the hg38 genome. To ensure realistic correlation between gene lengths and read counts, simulated gene counts were coupled to gene lengths. Specifically, gene counts and gene lengths were independently ranked, and lengths were then assigned to genes by matching these rank orders. Gaussian noise was added to the rank mapping to control the strength of the association, thereby inducing a strong but imperfect correlation between gene count and length.

Each gene was then divided into alternating simulated exonic and intronic features. The number of features created for each gene was designed such that expected exonic and intronic feature lengths were approximately 200 bp and 800 bp, respectively. Feature lengths were generated using a multinomial model, with feature specific probabilities drawn from a gamma distribution to create variable feature lengths. Counts were then distributed between exonic and intronic features according to a binomial model with target proportions of 40% and 60%, respectively. Each simulated feature was then further divided into fixed width windows (100 bp) and counts were distributed across windows using window weights drawn from a gamma distribution. This gamma distribution included an exonic and intronic specific peakiness parameter that controlled how evenly counts were distributed across all windows. Cross-sample noise drawn from a normal distribution was added to window weights to reflect realistic variance in window count distribution.

At this stage, four replicates were designated as IN samples and four as IP samples. IN replicates were left unmodified. Within the IP samples, 20,000 simulated genomic features were randomly selected and designated as bound features. For each bound feature *j*, a baseline binding strength, *κ_j_*, was drawn from a shifted gamma distribution, ensuring a strictly positive binding effect while allowing its magnitude to vary across features. A feature-specific log fold change, *ζ_j_*, was drawn from a normal distribution and exponentiated to obtain a strictly positive multiplicative treatment effect, exp(*ζ_j_*). Cross-replicate variation was then introduced by perturbing each feature’s treatment effect with noise drawn independently for each replicate. For each bound feature, the expected number of binding-associated reads in each IP replicate was calculated as the product of the feature-level count and the corresponding binding multiplier. Additional cross-replicate pulldown variability was introduced using a weak log-normal noise term. The resulting values were rounded to integers to generate pulldown counts. These counts were then added to a single window within the corresponding feature. The true binding strength and treatment-effect parameters for each feature were retained for the calculation of quality metrics.

All simulation parameters were estimated from the NONO eCLIP dataset (fig:figure5, Fig. S2, and Fig. S3). The complete simulation code, parameter values, and implementation details are available at https://github.com/YeoLab/flipper_simulation.

## 3 Results

### 3.1 Overview of Flipper and analysis strategy

Flipper addresses two central challenges in differential eCLIP analysis: normalization of IP signal across conditions and control for changes in RNA substrate availability. After peak calling and preprocessing, only regions showing significant IP enrichment in at least one condition are retained for analysis (Fig. 1A). This ensures that only relevant regions with adequate basal binding levels are analyzed by Flipper. Then, window-level IN counts are replaced with feature-level IN counts (INf) for later use in RNA substrate control. IP and IN data are normalized separately, with IP normalization performed in two steps to account for enriched and background regions (Fig. 1B). A DESeq2 framework then jointly models assay type, experimental condition, and their interaction (Fig. 1C), identifying sites where changes in IP signal cannot be explained by corresponding changes in RNA substrate. Significant sites are reported genome wide and subjected to downstream analysis and visualization.

### 3.2 Benchmarking differential binding specificity on eCLIP data

Both NONO and DDX42 datasets include vehicle and inactive compound control groups, enabling direct assessment of specificity and reproducibility in differential eCLIP analysis. Minimal differential binding is expected between the vehicle (DMSO) and inactive compound control groups (S-SKBG-1 or WX-02-43), while substantial overlap in sites is expected when comparing each control to the active treatment condition (R-SKBG-1 or WX-02-23).

Differential analysis with Flipper detected no significant binding changes between the vehicle and inactive compound groups in the NONO dataset (Fig. 2A). In contrast, both vehicle vs active and inactive vs active comparisons exhibit a noticeable shift toward increased binding, consistent with prior results (Kathman et al., 2023b).

**Figure 2:**
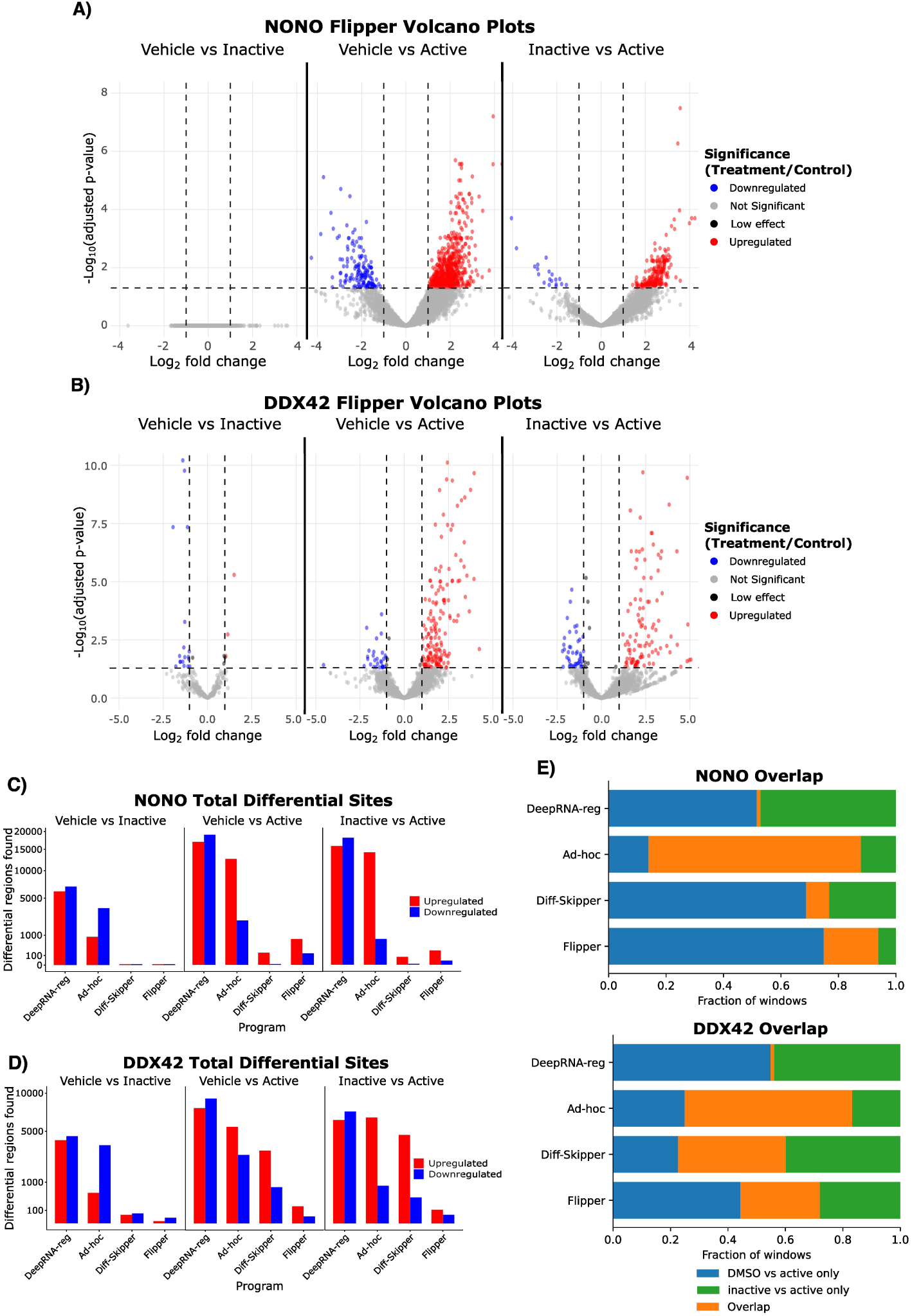
Method comparison for NONO and DDX42 eCLIP datasets. A-B) Volcano plots showing differential binding results for all comparisons in the NONO and DDX42 datasets, respectively. Minimal significant differences are observed between vehicle (DMSO) and the inactive compounds (S-SKBG-1 and WX-02-43), consistent with their expected lack of biological effect. C-D) Grouped bar plots summarizing the number of significantly up and downregulated sites identified by each differential binding method. Upregulated sites correspond to increased binding following treatment, whereas downregulated sites indicate decreased binding. E) Overlap bar plots showing concordance of differential sites identified in the DMSO vs active compound and inactive compound vs active compound comparisons. Alt text: Graphs depicting benchmarking results using real eCLIP datasets. Subfigures labelled a–e show statistical analyses and comparisons of the number and agreement of differential sites identified across programs.

In the DDX42 dataset, fewer than 1% of tested regions were differentially bound between the vehicle and inactive compound conditions, consistent with a low false positive rate under null conditions (Fig. 2B) (Supplementary Table 1). In contrast, a comparison of the vehicle and active compound conditions revealed a modest shift toward increased binding, although the total number of differential sites identified was unexpectedly low given the results reported by Lazear et al. (2023a). This discrepancy is discussed in section 3.3. However, for the purposes of initial validation, the key observation is that substantially fewer differential sites were identified between the control and inactive conditions than between the control and active conditions.

Across methods, Flipper and Diff-Skipper consistently identify few or no sites in vehicle vs inactive comparisons (Fig. 2C-D). In contrast, the ad-hoc method and DeepRNA-reg identify substantial numbers of differential sites under null conditions. Furthermore, while most programs recover the expected net increase in NONO binding following treatment, DeepRNA-reg identifies more sites with decreased binding than increased binding (Fig. 2C).

Consistency was assessed by examining overlap between sites identified in vehicle vs active and inactive vs active comparisons. Flipper and Diff-Skipper showed similar overlap, with Diff-Skipper showing slightly greater overlap in the DDX42 dataset and Flipper in the NONO dataset (Fig. 2E) (Supplementary Table 1). The ad-hoc method exhibited the highest overlap overall but also identified a large fraction of tested sites as significant, complicating interpretation of overlap as evidence of consistent site-specific detection. DeepRNA-reg showed near-zero overlap between contrasts (Fig. 2E).

dCLIP identified hundreds of thousands to millions of regions as differentially bound even in vehicle vs inactive compound comparisons (Supplementary Table 1). Given the scale of these results, their inclusion in Figure 2 would have obscured meaningful comparison with other methods. DeepRNA-reg also produced large numbers of differential calls under null contrasts and showed minimal similarity between treatment comparisons. This behavior makes it difficult to perform interpretable, meaningful comparisons between dCLIP, DeepRNA-reg and other programs. As such, subsequent benchmarking was restricted to Flipper, the ad-hoc method, and Diff-Skipper.

### 3.3 Flipper accounts for differential splicing effects

The unexpectedly small number of differential binding sites identified in the DDX42 dataset prompted further investigation into how substrate availability controls affect Flipper results. Flipper provides two options for RNA substrate control: the default feature-level input control (INf) and a gene-level input control (INg). Because INg estimates RNA substrate availability using gene-wide counts, it generally provides greater statistical power than INf, which uses fewer reads from the local feature. Consequently, INf is expected to have lower sensitivity. This behavior is observed in the PUF60 dataset, where INg and INf produce broadly similar results with only a modest reduction in the number of significant sites identified by INf (Fig. 3A).

**Figure 3:**
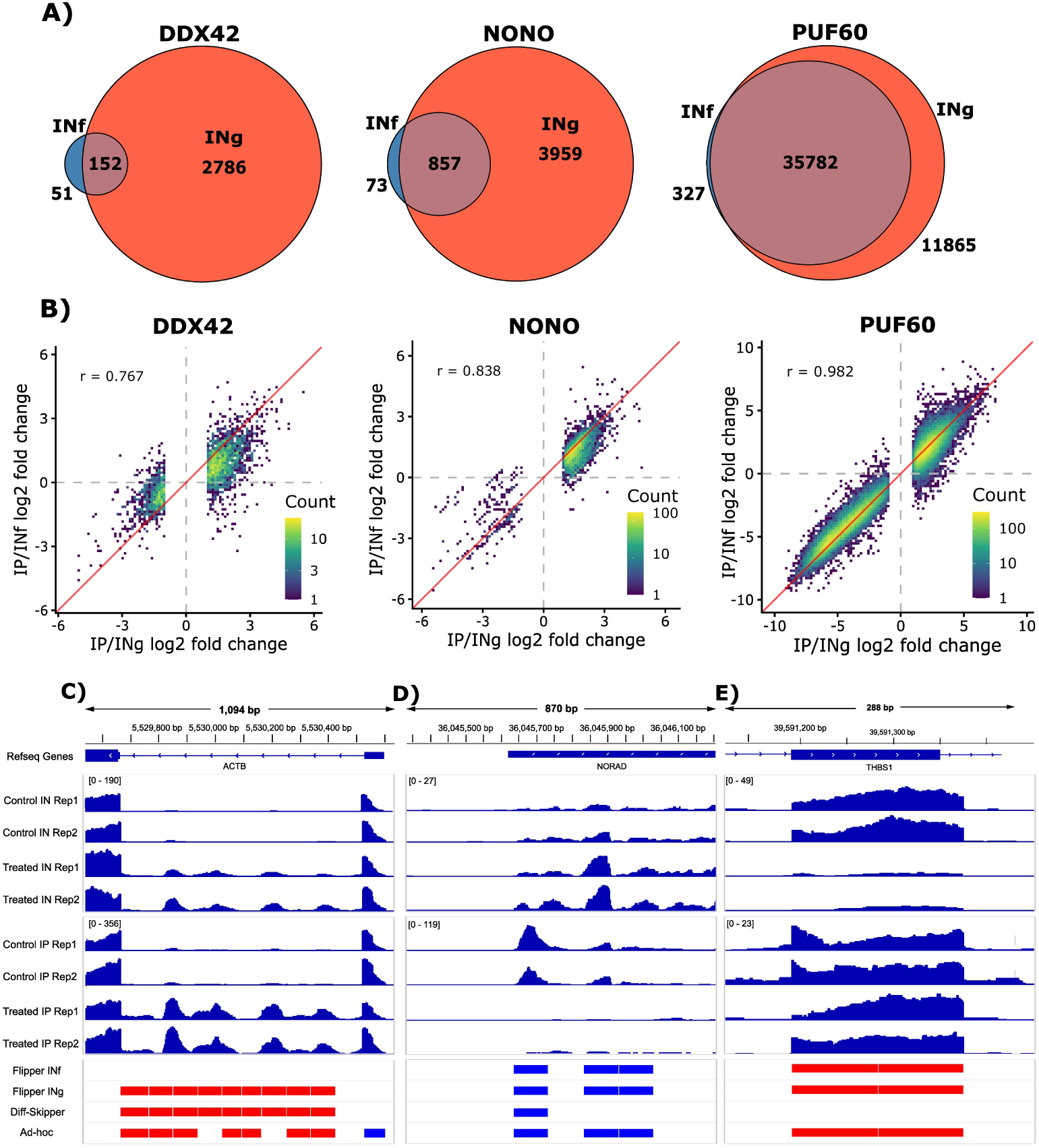
Comparison of Flipper results using feature-wise IN (INf) and gene-wise IN (INg) RNA substrate availability control. A) Venn diagrams showing overlap between differential sites identified using either method. B) Scatter plots comparing IP/INf and IP/INg log2 fold changes for all sites identified as significant by either method. C-E) Integrated Genomics Viewer (IGV) scaled coverage tracks. Counts are scaled using size factors created by Flipper’s hierarchical normalization method. Horizontal red and blue bars indicate significantly up and downregulated differential sites, respectively. C) Region identified as having differential DDX42 binding activity by all methods except the INf version of Flipper. Region shows similar increases in both IP and IN signal consistent with intron retention rather than differential binding. D) Region identified as having differential PUF60 binding activity by all methods. Region shows a substantial decrease in IP signal with relatively stable IN coverage across conditions. E) Region showing differential PUF60 binding activity by all methods except Diff-Skipper. Region shows stable IP signal accompanied by a substantial decrease in local IN signal, consistent with increased relative binding following treatment. Alt text: Graphs depicting the importance of controlling for splicing effects using feature-wise input during differential eCLIP analysis. Subfigures labelled a–e compare results across input control methods and show genomic tracks highlighting affected regions.

In contrast, INg and INf produced substantially greater differences in the NONO and DDX42 datasets than expected from reduced statistical power alone (Fig. 3A), suggesting an additional source of disagreement. To investigate, log_2_ fold changes obtained using INg and INf were compared across all sites identified as significant in either mode (Fig. 3B). In the PUF60 dataset, fold changes remained strongly correlated between modes, with only minor disagreement at a small subset of sites. In contrast, agreement was much lower in the NONO dataset, while the DDX42 dataset showed widespread disagreement between modes.

Inspecting the genomic regions contributing to this disagreement indicated that alternative splicing contributed to the differences between INf and INg. For example, in the DDX42 dataset, IP and IN signals increased by similar amounts at an ACTB intron following treatment, suggesting increased RNA substrate abundance rather than increased relative binding (Fig. 3C). INf increased substantially, while INg remained stable, indicating a local rather than gene-wide change in RNA substrate levels consistent with intron retention (Supplementary Table 2). IRFinder further supported intron retention at this site, showing a 5.66-fold increase in IR ratio following treatment (Supplementary Table 3). Flipper correctly did not classify the site as differentially bound when in INf mode, whereas Flipper in INg mode and all other methods tested classified it as differentially bound. This distinction is particularly important in the DDX42 dataset because the treatment was specifically designed to perturb splicing. Additional examples are shown in Supplementary Figure 4.

(Fig. 3D) shows an example from the PUF60 dataset in which all differential binding methods identify a substantial loss of binding following treatment. This loss was driven almost entirely by a change in IP signal, while local INf and gene-level INg signals remained stable across conditions.

(Fig. 3E) shows an example from the PUF60 dataset in which IP signal remains stable across conditions, while local IN signal decreases substantially following treatment. This implies increased binding relative to substrate availability. Because Diff-Skipper relies exclusively on IP signal, this site is missed. In contrast, Flipper detects the change because it specifically addresses changes in RNA substrate abundance. This illustrates that appropriate IN control can facilitate the detection of meaningful changes in binding.

Overall, these results suggest that feature-level input control is necessary to distinguish true differential binding from changes driven by alternative splicing or RNA expression. While the DDX42 dataset still exhibits some evidence of differential binding following treatment, altered DDX42 binding does not fully explain the substantial splicing effects of WX-02-23. This may motivate additional experiments exploring alternative hypotheses, such as WX-02-23 modulating DDX42’s interaction with spliceosomal proteins.

### 3.4 Flipper visualizations enable downstream interpretation of differential binding

dCLIP, DeepRNA-reg, and the ad-hoc method produce relatively simple outputs, typically consisting of tables of increased and decreased binding sites, with dCLIP also providing BED files of differential regions. Repurposed peak callers such as Diff-Skipper generate more visualizations, but these are designed for peak detection rather than differential analysis. In contrast, Flipper performs additional downstream analyses and produces visualizations designed to facilitate interpretation of differential binding results. The following examples are drawn from the PUF60 dataset comparing wild type PUF60 with the L140P mutant, which has previously been shown to exhibit altered RNA binding activity (Tankka et al., 2025a).

Flipper generates summary statistics for each gene containing at least one significantly differentially bound site. For each gene, Flipper reports Fisher combined p-values and total (summed) log-fold change across all significant sites, emphasizing genes with large aggregate changes. Flipper also provides the minimum p-value per gene, highlighting genes with individually strong sites. Gene-level summaries and associated visualizations are shown in Fig. S5.

Flipper generates a volcano plot summarizing effect sizes (Fig. 4A) and a diverging bar plot showing the distribution of up and downregulated binding sites across genomic feature types (Fig. 4B). Consistent with prior reports, Flipper analysis finds that the L140P mutation is associated with decreased binding at canonical intronic PUF60 sites. Notably, we also observe an increase in binding within coding sequence (CDS) regions. While *in vitro* studies suggest that the mutation reduces binding to canonical UC rich motifs (Kralovicova et al., 2020), its broader in vivo effects remain less well characterized. Motif analysis using HOMER (Heinz et al., 2010) further supports this distinction: downregulated sites are enriched for the known UC rich PUF60 binding motif, whereas upregulated sites lack a clear motif (Fig. 4C).

**Figure 4:**
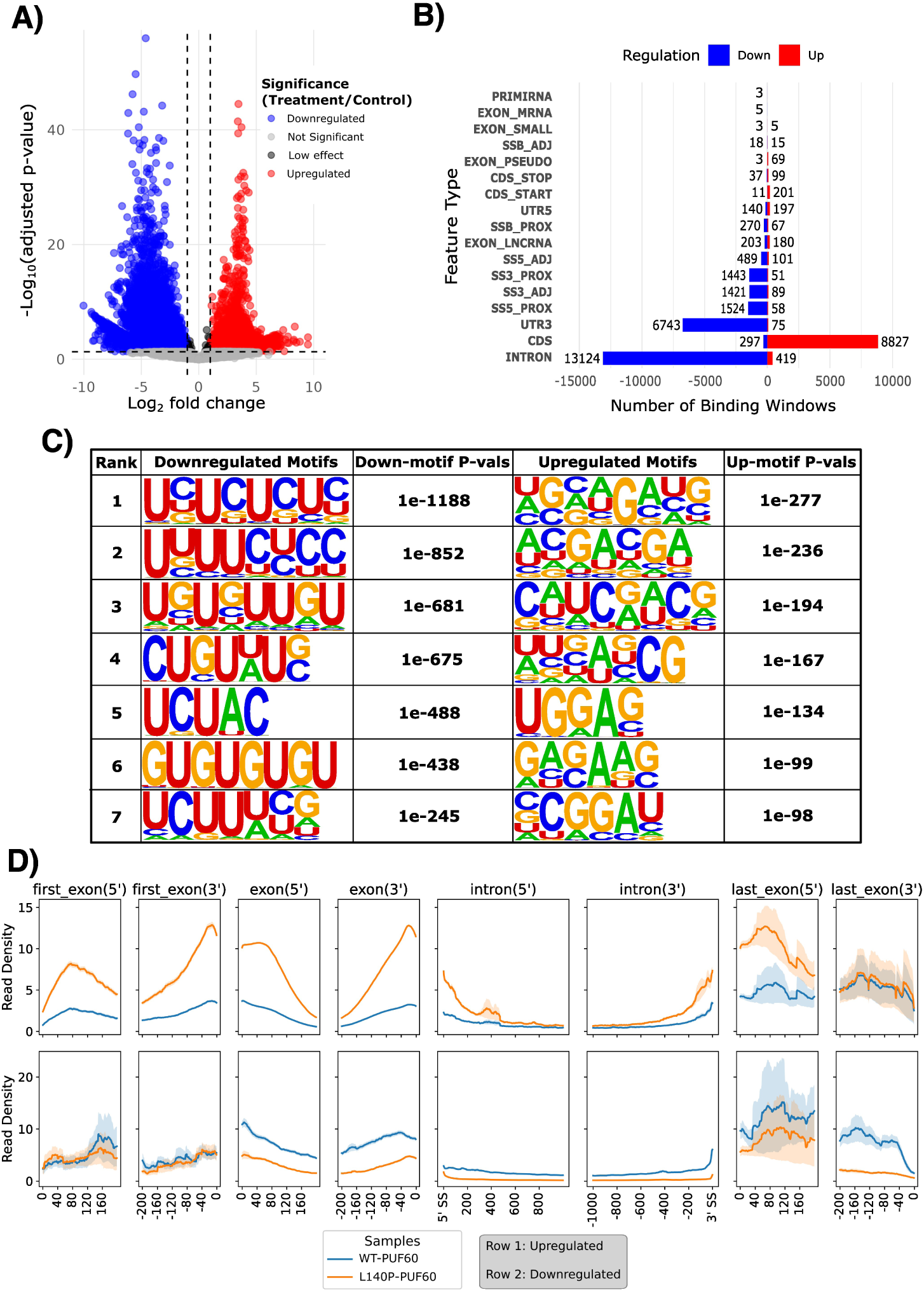
Flipper generated visualizations for the PUF60 eCLIP dataset. A) Volcano plot showing widespread increases and decreases in binding following the L140P mutation. B) Diverging bar plot summarizing genomic feature enrichment, highlighting decreased binding within intronic and 3*^′^* UTR regions and increased binding within coding sequences (CDS). C) HOMER motif analysis of up and downregulated windows, with several UC rich motifs enriched among downregulated sites. D) Metadensity plots showing relative read density across genomic regions containing significantly up or downregulated binding sites in control and treatment conditions. Read density is estimated using the relative information (RI) metric from the original metadensity program. As expected, regions with upregulated binding sites exhibit increased treatment read density, whereas regions with downregulated binding sites show the opposite trend. Alt text: Graphs depicting the visualization outputs generated by Flipper. Subfigures labelled a–d show volcano plots, feature type barplots, motif analysis results, and metadensity plots.

To further characterize this redistribution, Flipper adapts the metadensity framework (Her et al., 2022) to visualize eCLIP signal across genomic features containing differential sites. These metadensity plots show that the L140P mutant exhibits increased read density near both 5*^′^* and 3*^′^* splice sites in regions containing upregulated binding sites, particularly within exonic regions (Fig. 4D, top row). A similar trend is observed in features that contain downregulated binding sites, although the change is subtler (Fig. 4D, bottom row).

Together, these visualizations indicate that, in addition to reducing binding at canonical PUF60 sites, the L140P mutation is associated with a redistribution of binding toward exonic regions flanking splice sites. While this finding arose from an exogenous expression system and the mechanistic basis of this shift remains to be determined, it illustrates how Flipper enables detection of broader changes in binding architecture.

### 3.5 Benchmarking on simulated datasets

Although analysis of real datasets provides insight into the behavior of each method, quantitative performance metrics require access to known ground truth. To directly assess performance under controlled conditions, we evaluated Flipper, the ad-hoc method, and Diff-Skipper using simulated data.

Simulated eCLIP count data were generated via a three-step process. First, simulated genelevel counts were drawn from a negative binomial distribution (Fig. 5A). Next, these gene-level counts were divided into simulated intronic and exonic features before being further divided into individual genomic windows (Fig. 5B). Finally, individual features were designated as bound and counts at a specific window within the feature were enriched as a function of the number of reads attributed to the feature and a pulldown parameter (*κ*) drawn from a gamma distribution (Fig. 5C). Additional simulation details are provided in Section 2.7.

**Figure 5:**
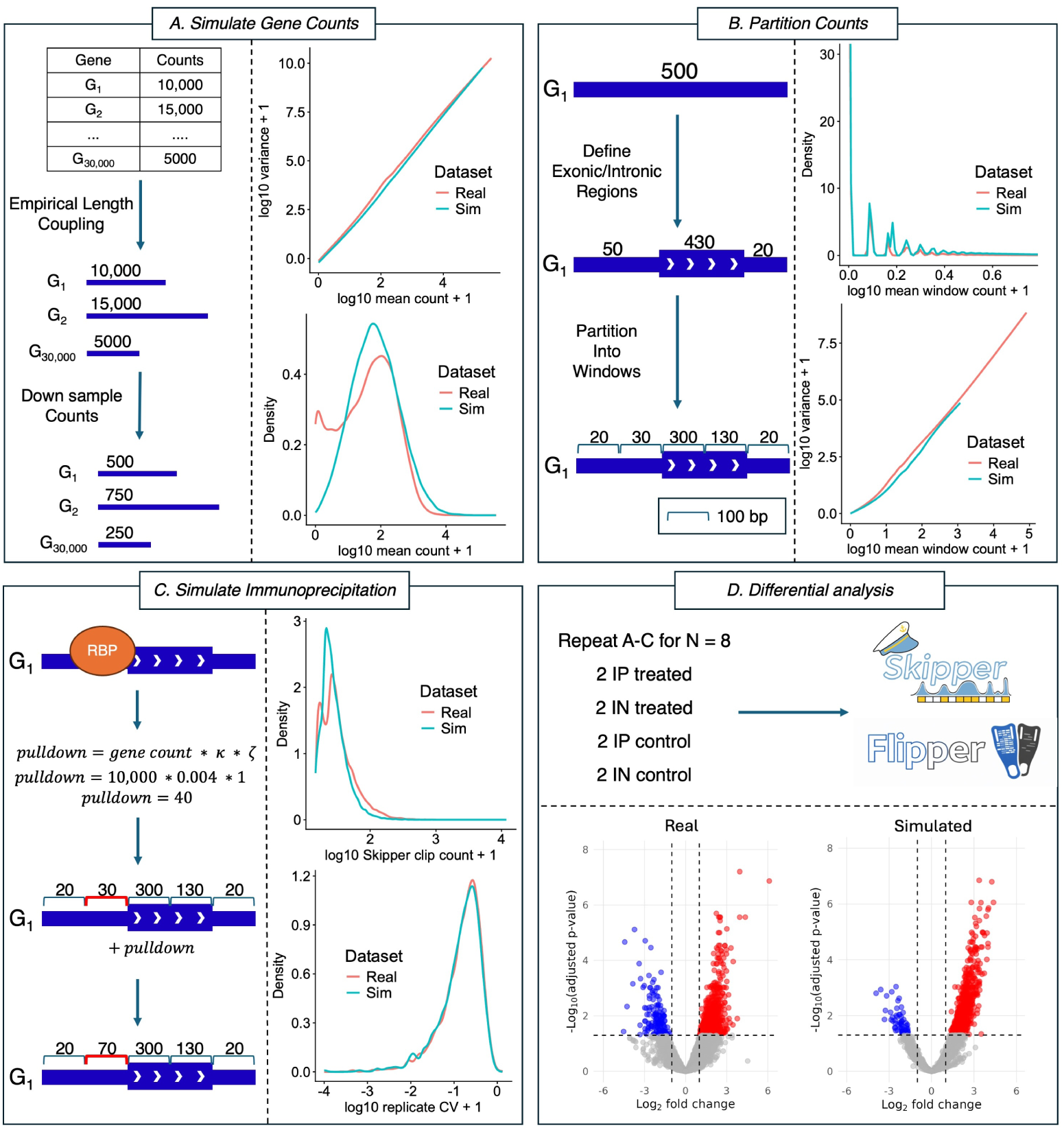
Overview of eCLIP simulation pipeline with comparisons between real DDX42 eCLIP data and simulated eCLIP data. Each panel shows a schematic of a major simulation step and comparisons between real and simulated data. A) Initial gene counts are generated from a negative binomial distribution and coupled to empirical gene lengths before being downsampled to reflect the sparsity of eCLIP data. Simulated gene-level counts exhibit count distributions and mean-variance relationships that closely resemble those observed in real data. B) Counts are divided into high and low read depth regions and then further partitioned into 100 bp windows. High similarity is observed between real and simulated window-level counts. C) Immunoprecipitation is simulated by adding a pulldown value to a specific window for each bound gene. The count and cumulative variance (CV) distributions for significant binding windows identified by Skipper in both the real and simulated data are very similar. D) The simulation is repeated to generate IP and IN data for both treatment groups, and these data are then subjected to differential binding analysis. Volcano plots show similar distributions between real and simulated data. Alt text: Diagram depicting the main steps of the eCLIP simulation pipeline. The workflow is split into four subpanels labelled a–d, covering gene-level count simulation, window generation, simulated binding enrichment, and differential analysis.

Each simulated experiment consisted of eight replicates: four IN and four IP samples distributed across control and treatment groups. In the treatment condition, the simulated pulldown parameter at each binding site was modulated by a treatment specific binding effect *ζ*. Expression levels were either constant between control and treatment groups (mean *β* = 0, standard deviation *β* = 0) or allowed to vary under treatment (mean *β* = 0, standard deviation *β* = 2). All simulation parameters, including *ζ*, sources of variability, and read depth were estimated from the NONO dataset (fig:figure5, Fig. S2, and Fig. S3).

The performance of each method was evaluated using sensitivity and precision. Sensitivity was defined as the fraction of true differential binding sites correctly identified, while precision was defined as the fraction of sites called significant that were truly differential. True differential sites were defined as those which had a *ζ* value above 1.5 or below 0.66 and which were identified by Skipper as binding sites in at least one experimental group. Five replicate simulations were performed for each treatment-parameter combination to assess the consistency of each method.

Figure 6 compares the sensitivity and precision between methods with and without expression modulation by the treatment. Flipper’s sensitivity hovers slightly above 20%, while precision remains consistently high at around 90%. Although Flipper’s sensitivity is modest, this likely reflects the inherent sparsity and variability of eCLIP count data, where true differential binding events may be difficult to detect without inflating false positives. The consistently high precision suggests that this tradeoff favors specificity in noisy experimental settings. Notably, Flipper’s performance is largely unaffected by treatment induced expression changes, demonstrating effective adjustment for expression effects.

**Figure 6:**
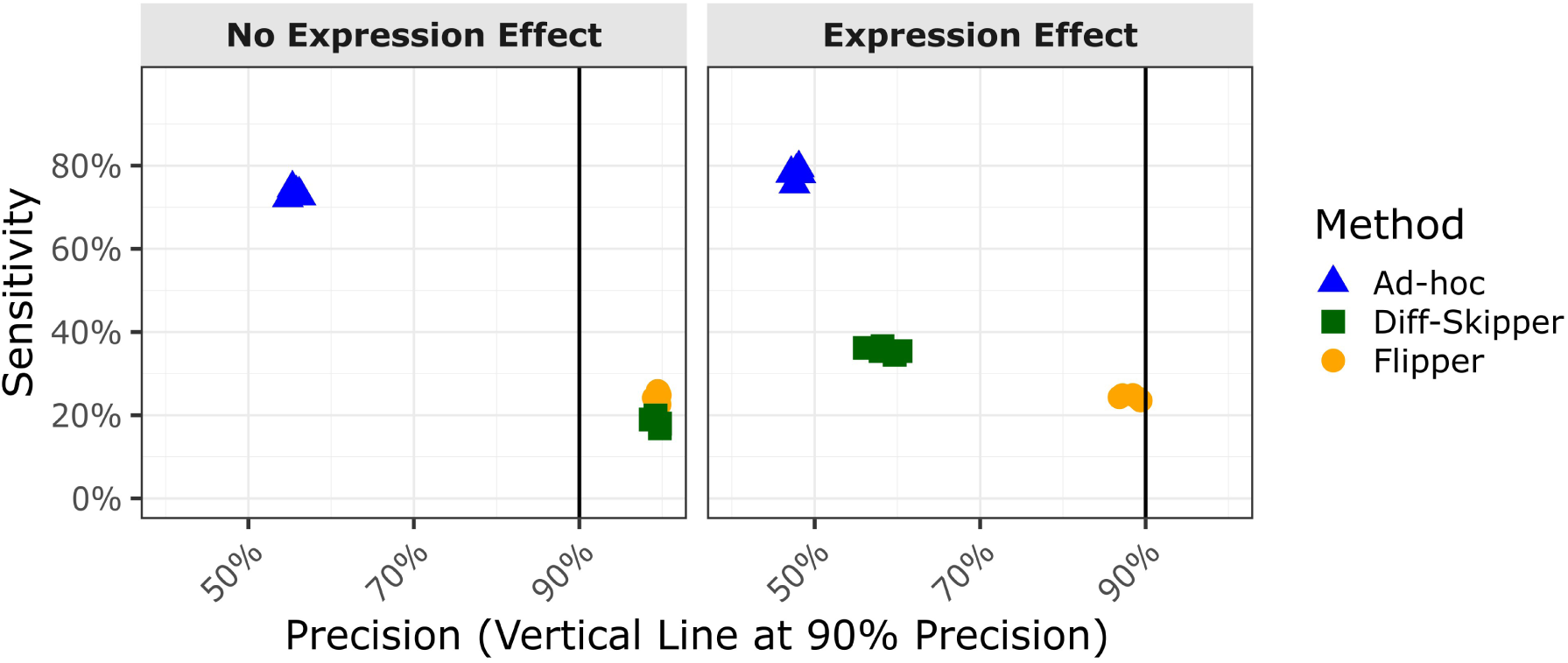
Sensitivity and precision of differential binding methods across simulated combinations of binding and expression treatment effects. Flipper performs comparably to or modestly better than alternative methods when expression effects are absent, and substantially outperforms other methods when expression changes are present, reflecting its ability to distinguish binding specific shifts from expression driven differences. Alt text: Graph comparing performance metrics across programs on simulated eCLIP data. Results show Flipper maintains high sensitivity and precision across the range of tested simulation parameters.

The ad-hoc method exhibits relatively stable performance across conditions, with sensitivity around 80% and precision around 50%, consistently yielding the lowest precision among the methods evaluated. In the absence of expression effects, Diff-Skipper follows a trend similar to Flipper, showing similar precision with slightly lower sensitivity. However, when expression changes are introduced, its precision declines sharply to approximately 60%, barely outperforming the ad-hoc method.

## 4 Discussion

Differential RBP binding analysis following treatment or mutation is an increasingly important component of both drug development and disease research. Despite this growing demand, existing approaches remain limited in their ability to address the unique statistical and experimental challenges inherent to using CLIP data for differential analysis. Here, we introduce Flipper, a method designed to address key challenges in differential RBP binding analysis, including control for RNA substrate availability and normalization strategies that account for signal to noise variation across samples. By leveraging the DESeq2 framework in conjunction with feature-level IN adjustments, Flipper consistently outperforms prior differential methods across both real and simulated datasets while also enabling expanded downstream analyses.

When evaluated on real datasets, Flipper exhibited low false positive rates in control to control comparisons and produced highly concordant results across control to treatment analyses. Furthermore, Flipper recapitulated previously reported biological findings, including increased binding in the NONO dataset and reduced binding to UC rich regions in the PUF60 dataset. Although other differential binding programs produced comparable differential site counts, examination of individual loci indicated that many were driven by RNA substrate changes rather than binding effects.

Analysis of simulated data further underscored this limitation. In the presence of treatment-induced changes in RNA expression, Flipper maintained substantially higher precision than the alternative methods, indicating greater robustness to expression-driven false positives. Although this higher precision was accompanied by lower sensitivity, the resulting tradeoff reflects a deliberate choice to favor reliable discovery in sparse, high-variance eCLIP datasets.

Notably, the published differential CLIP methods dCLIP and DeepRNA-reg did not perform as expected when evaluated on real eCLIP datasets. For dCLIP, this behavior may reflect a failure to satisfy its core assumption that global binding levels remain stable across conditions. Many treatments induce net increases or decreases in RBP binding, undermining this assumption. For DeepRNA-reg, one possible explanation is limited generalizability beyond its original training context. DeepRNA-reg was trained on a single HITS-CLIP dataset for the mouse microRNA mir-155 (Loeb et al., 2012) and subsequently evaluated on related microRNA CLIP datasets. Because microRNA binding events typically generate narrower peaks than those observed for most RBPs (Sekhon et al., 2025), this model may struggle with the broader peaks observed in many eCLIP studies.

Several limitations of Flipper warrant consideration. Feature-level input controls have less statistical power for short or lowly expressed features because local estimates of RNA substrate availability become less reliable as coverage decreases. To address this limitation, Flipper also supports gene-level input control, which may be useful when studying low-coverage features or in experimental settings where splicing changes are expected to be minimal. A second limitation is that the hierarchical normalization strategy used by Flipper assumes that, on average, pulldown efficiency does not differ systematically between treatment groups. This assumption is difficult to validate, but it enables detection of global shifts in binding that would otherwise be obscured. More robust normalization may ultimately require incorporation of external spike-ins, similar to those developed for ChIP-seq (Patel et al., 2024). The development of spike-in strategies for eCLIP represents an important future direction for improving differential RBP binding analysis.

Despite these limitations, our results demonstrate that Flipper provides a robust and principled framework for differential RBP binding analysis. By integrating substrate availability aware modeling with hierarchical normalization and detailed downstream analysis, Flipper enables rigorous identification and contextualization of condition specific changes in RBP binding. As differential eCLIP experiments continue to become more common, pipelines that explicitly disentangle binding and substrate effects will be essential for accurate biological inference.

## 5 Competing interests

No competing interest is declared.

## 6 Acknowledgements

We thank Summer R. Fair, Hema Kopalle, Nadav Wallis, and Brian Yee for their valuable discussions.

OpenAI’s ChatGPT (GPT-5.5) large language model was used as an aid to correct written text, to assist in code writing, and to generate Flipper’s logo. The authors are responsible for all scientific content, analyses, code, and manuscript text and take full responsibility for the accuracy of the work.

## 7 Author contributions statement

Keegan Flanagan (Conceptualization [lead], Data Curation [lead], Formal analysis [lead], Methodology [lead], Software [lead], Validation [lead], Visualization [lead], Writing-original draft [lead]), Shuhao Xu (Conceptualization [supporting], Data Curation [supporting], Formal analysis [supporting], Software [supporting], visualization [supporting], Writing-review and editing [equal]), Gene Yeo (Conceptualization [supporting], Funding acquisition [lead], Project administration [lead], Resources [lead], Supervision [lead], Writing-review and editing [equal])

## 8 Funding

This work was supported by the National Institutes of Health [R01 HG004659,U24HG009889] and the Natural Sciences and Engineering Research Council of Canada Post Graduate Scholarship-Doctoral program.

## 9 Data availability

The Flipper and Skipper software packages are freely available under the UC license on GitHub (https://github.com/YeoLab/flipper and https://github.com/YeoLab/skipper, respectively). A versioned archive of Flipper is also available on Zenodo (https://doi.org/10.5281/zenodo.21136504). Additional benchmarking and evaluation scripts used to generate the manuscript figures are available at https://github.com/YeoLab/flipper/tree/main/manuscript_figures. All analysis scripts, configuration files, and benchmarking workflows used for the simulation analyses are available at https://github.com/YeoLab/flipper_simulation.

The GRCh38 reference genome and gene annotations were obtained from GENCODE (https://ftp.ebi.ac.uk/pub/databases/gencode/Gencode_human/release_49/gencode.v49.basic.annotation.gff3.gz). eCLIP data for the NONO, DDX42, and PUF60 RNA-binding proteins are available from the Gene Expression Omnibus (GEO; https://www.ncbi.nlm.nih.gov/geo/) under accession numbers GSE198212, GSE220185, and GSE280899, respectively.

## Notes

### Competing Interest Statement

The authors have declared no competing interest.

### Summary of Updates

Final pre-submission update. Major changes include a switch from INg to INf for the default input control and various simulation updates.

