## Supplemental Info for "Flipper: An advanced framework for identifying differential RNA binding behavior with eCLIP data"

**Table 1.** Summary of differential binding calls and overlap across methods, datasets, and contrasts.

| Program | Dataset | Comparison | Total sites tested | Significant sites | Overlapping sites | Jaccard index |
|---|---|---|---|---|---|---|
| Diff-Skipper | NONO | DMSO vs S-SKBG-1 | Transcriptome | 1 | NA | NA |
| | | DMSO vs R-SKBG-1 | Transcriptome | 172 | 18 | 0.0804 |
| | | S-SKBG-1 vs R-SKBG-1 | Transcriptome | 70 | | |
| | DDX42 | DMSO vs WX-02-43 | Transcriptome | 102 | NA | NA |
| | | DMSO vs WX-02-23 | Transcriptome | 3907 | 2433 | 0.3743 |
| | | WX-02-43 vs WX-02-23 | Transcriptome | 5026 | | |
| Adhoc Method | NONO | DMSO vs S-SKBG-1 | 6733 | 4528 | NA | NA |
| | | DMSO vs R-SKBG-1 | 18795 | 15226 | 12811 | 0.7389 |
| | | S-SKBG-1 vs R-SKBG-1 | 17271 | 14924 | | |
| | DDX42 | DMSO vs WX-02-43 | 4885 | 4150 | NA | NA |
| | | DMSO vs WX-02-23 | 9843 | 8281 | 5797 | 0.5828 |
| | | WX-02-43 vs WX-02-23 | 7907 | 7463 | | |
| DeepRNA-reg | NONO | DMSO vs S-SKBG-1 | 6733 | 12965* | NA | NA |
| | | DMSO vs R-SKBG-1 | 18795 | 36983* | 904 | 0.0129 |
| | | S-SKBG-1 vs R-SKBG-1 | 17271 | 33890* | | |
| | DDX42 | DMSO vs WX-02-43 | 4885 | 8561* | NA | NA |
| | | DMSO vs WX-02-23 | 9843 | 17047* | 367 | 0.0121 |
| | | WX-02-43 vs WX-02-23 | 7907 | 13715* | | |
| Flipper | NONO | DMSO vs S-SKBG-1 | 6733 | 0 | NA | NA |
| | | DMSO vs R-SKBG-1 | 18795 | 930 | 188 | 0.1897 |
| | | S-SKBG-1 vs R-SKBG-1 | 17271 | 249 | | |
| | DDX42 | DMSO vs WX-02-43 | 4885 | 22 | NA | NA |
| | | DMSO vs WX-02-23 | 9843 | 203 | 78 | 0.2766 |
| | | WX-02-43 vs WX-02-23 | 7907 | 157 | | |
| dCLIP | NONO | DMSO vs S-SKBG-1 | Transcriptome | 3167674 | NA | NA |
| | | DMSO vs R-SKBG-1 | Transcriptome | 3182495 | 2753744 | 0.8653 |
| | | S-SKBG-1 vs R-SKBG-1 | Transcriptome | 2753744 | | |
| | DDX42 | DMSO vs WX-02-43 | Transcriptome | 1004625 | NA | NA |
| | | DMSO vs WX-02-23 | Transcriptome | 1478820 | 1004625 | 0.6793 |
| | | WX-02-43 vs WX-02-23 | Transcriptome | 1216098 | | |

**Table 2.** Comparison of feature-level (INf) and gene-level (INg) input counts in control and treatment samples. Mean counts were calculated across the two replicates from each condition. Fold change was calculated as the mean treatment count divided by the mean control count. These measurements correspond to the four examples shown in Supplementary Figure 4.

| Gene | Input | Treatment 1 | Treatment 2 | Control 1 | Control 2 | Mean treatment | Mean control | Fold change |
|---|---|---|---|---|---|---|---|---|
| ACTB | INf | 231 | 238 | 60 | 51 | 234.5 | 55.5 | 4.23 |
| | INg | 1504 | 1203 | 1372 | 1236 | 1353.5 | 1304.0 | 1.04 |
| HNRNPH1 | INf | 52 | 57 | 45 | 26 | 54.5 | 35.5 | 1.54 |
| | INg | 1672 | 1153 | 2538 | 2098 | 1412.5 | 2318.0 | 0.61 |
| TPT1 | INf | 82 | 66 | 24 | 11 | 74.0 | 17.5 | 4.23 |
| | INg | 796 | 596 | 942 | 761 | 696.0 | 851.5 | 0.82 |
| SRRM2 | INf | 22 | 10 | 4 | 2 | 16.0 | 3.0 | 5.33 |
| | INg | 2718 | 1954 | 3607 | 2582 | 2336.0 | 3094.5 | 0.75 |

**Table 3.** IRFinder quantification of intron retention in control and treatment samples for four representative introns. Intron depth represents the mean number of intronic reads per base pair. Splice left and Splice right indicate the numbers of splice junction reads overlapping the 5′ and 3′ flanking exons, respectively. IR ratio was calculated as $\mathrm{IRratio} = \mathrm{IntronDepth}/[\max(\mathrm{SpliceLeft}, \mathrm{SpliceRight}) + \mathrm{IntronDepth}]$. Mean IR ratios were calculated across two replicates per condition, and IR fold change was calculated as the mean treatment IR ratio divided by the mean control IR ratio. These measurements correspond to the four examples shown in Supplementary Figure 4.

| Intron | Gene | Sample | Intron depth | Splice left | Splice right | IR ratio | Mean IR ratio | IR fold change |
|---|---|---|---|---|---|---|---|---|
| chr7:5529663–5530523 | ACTB | Control 1 | 2.81 | 147 | 147 | 0.0188 | 0.0190 | 5.66 |
| | | Control 2 | 2.60 | 132 | 133 | 0.0192 | | |
| | | Treatment 1 | 11.38 | 111 | 111 | 0.0930 | 0.1073 | |
| | | Treatment 2 | 9.97 | 72 | 72 | 0.1217 | | |
| chr5:179623164–179623594 | HNRNPH1 | Control 1 | 4.73 | 7 | 4 | 0.4030 | 0.3663 | 1.80 |
| | | Control 2 | 2.46 | 5 | 4 | 0.3296 | | |
| | | Treatment 1 | 5.23 | 3 | 3 | 0.6356 | 0.6608 | |
| | | Treatment 2 | 4.37 | 2 | 2 | 0.6860 | | |
| chr13:45340785–45341041 | TPT1 | Control 1 | 4.69 | 21 | 21 | 0.1826 | 0.1903 | 3.53 |
| | | Control 2 | 2.47 | 10 | 10 | 0.1981 | | |
| | | Treatment 1 | 18.19 | 11 | 12 | 0.6025 | 0.6713 | |
| | | Treatment 2 | 14.23 | 4 | 5 | 0.7400 | | |
| chr16:2759047–2759139 | SRRM2 | Control 1 | 1.00 | 35 | 35 | 0.0278 | 0.0139 | 15.13 |
| | | Control 2 | 0.00 | 22 | 22 | 0.0000 | | |
| | | Treatment 1 | 4.41 | 10 | 11 | 0.2863 | 0.2102 | |
| | | Treatment 2 | 2.32 | 12 | 15 | 0.1341 | | |

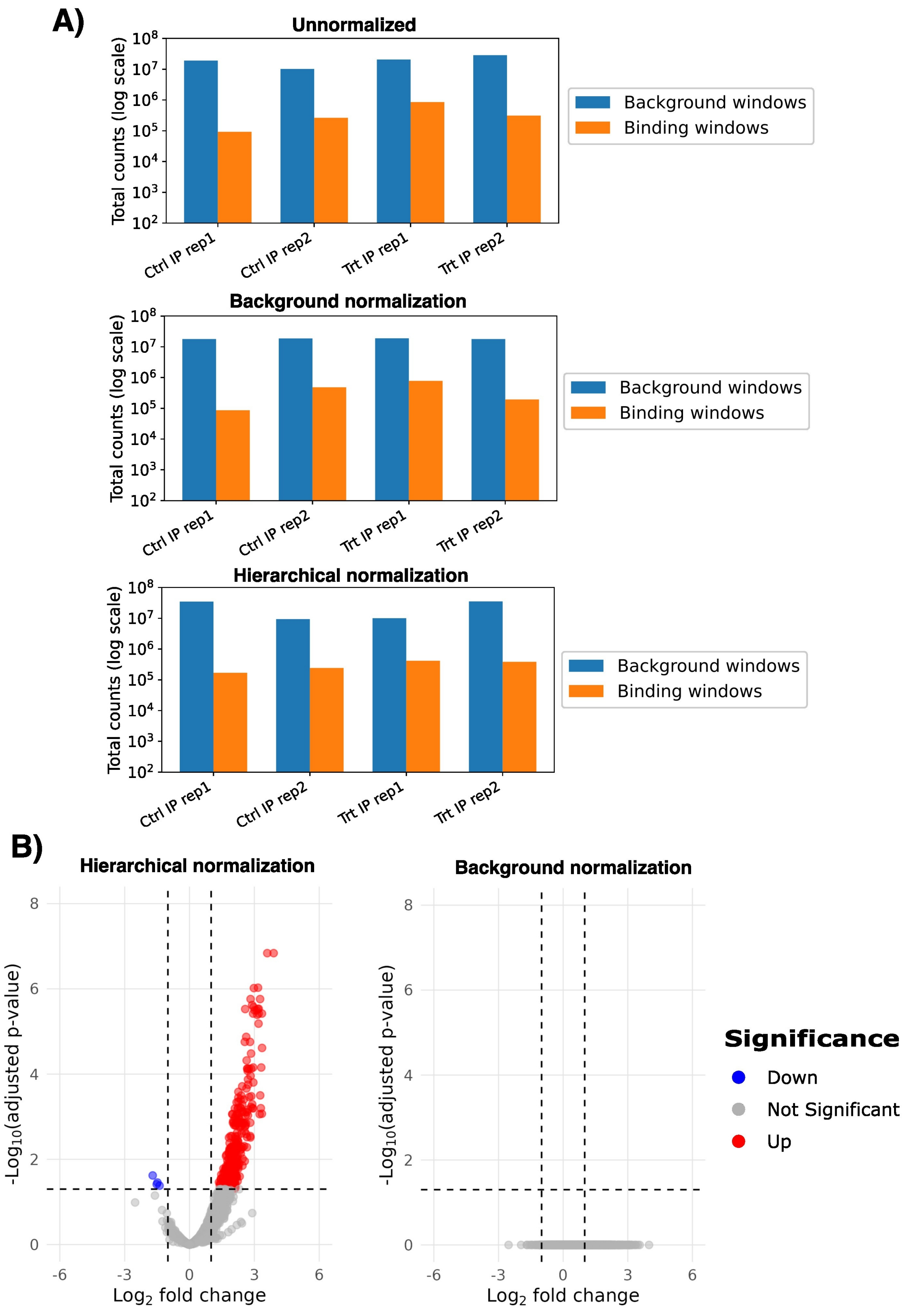


Fig. 1: Demonstration of the importance of hierarchical normalization under simulated signal to noise ratio shifts. A) Comparison of replicate-level read counts across normalization strategies. Top: unnormalized total read counts, where relative differences between replicates are reversed when comparing signal and background windows. Middle: traditional background based normalization, which amplifies replicate separation within binding windows. Bottom: hierarchical normalization, which restores comparable binding window counts across replicates while preserving appropriate control of overall sequencing depth between treatment groups. B) Volcano plots comparing differential binding results obtained using hierarchical versus background normalization. Increased replicate variance introduced by background normalization markedly reduces the number of significant windows detected. Note that a net increase in binding is expected from this simulation

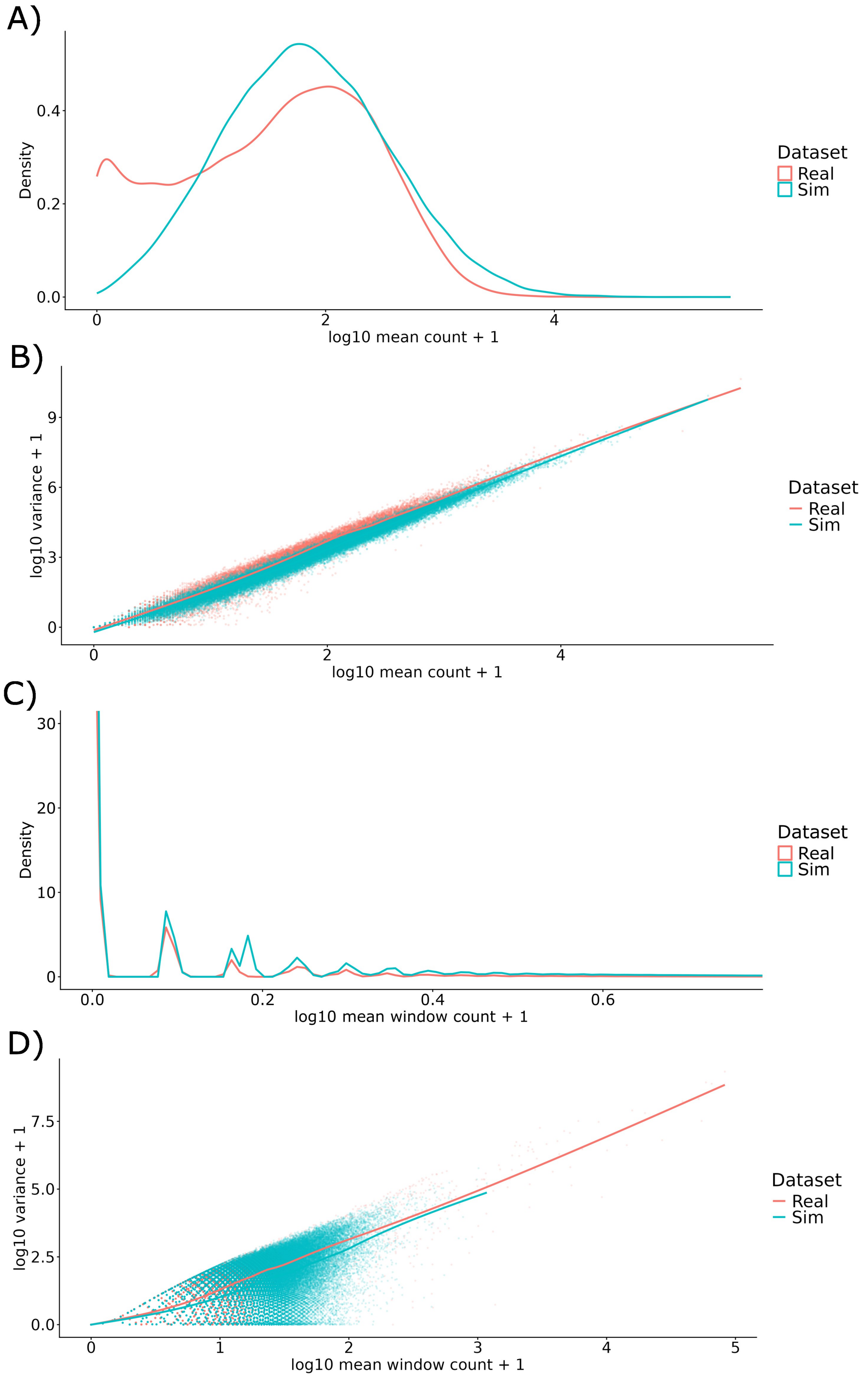


Fig. 2: Expanded comparisons between real and simulated data. This plot shows the same comparison between the NONO and simulated data shown in figure 5 panels A-B, but at an expanded scale and with visible points where appropriate. A) Gene-level count distributions for real and simulated data. Real and simulated distributions exhibit similar shapes, though real data clearly exhibits a higher number of un-expressed genes. B) Gene-level count mean-variance relationships for real and simulated data. Simulated data recapitulate variability between replicates. C) Window-level count distributions for real and simulated data. Real and simulated distributions exhibit similar shapes. D) Window-level count mean-variance relationships for real and simulated data. Simulated data recapitulate variability between replicates.

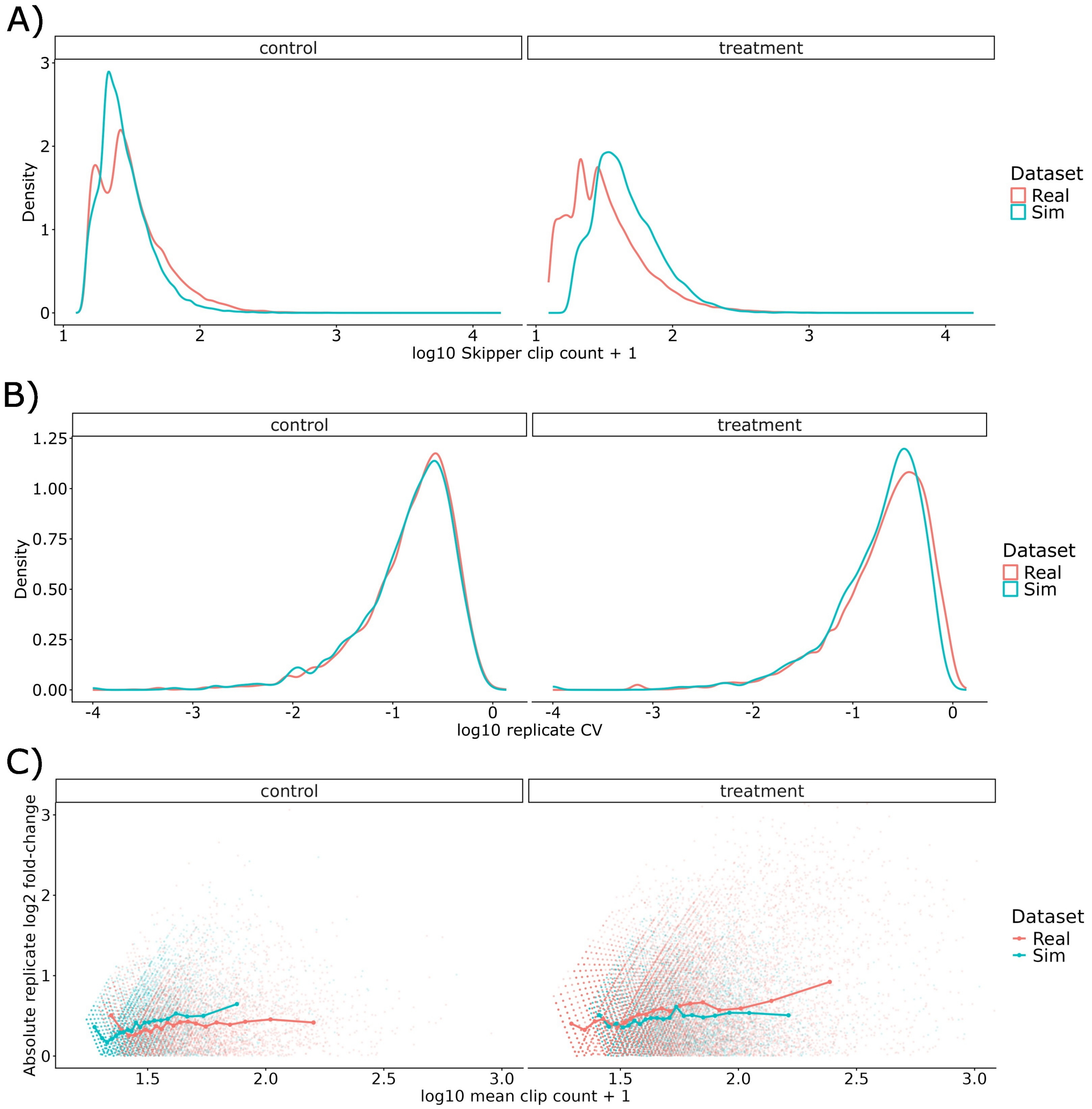


Fig. 3: Additional comparisons between real and simulated Skipper output. A) Count distributions for real and simulated data. B) Distributions of coefficient of variation (CV) for real and simulated data. C) Mean-variance relationships for real and simulated data. Replicate log2 fold changes represent differences between replicates within the same condition. Simulated data recapitulate variability between replicates.

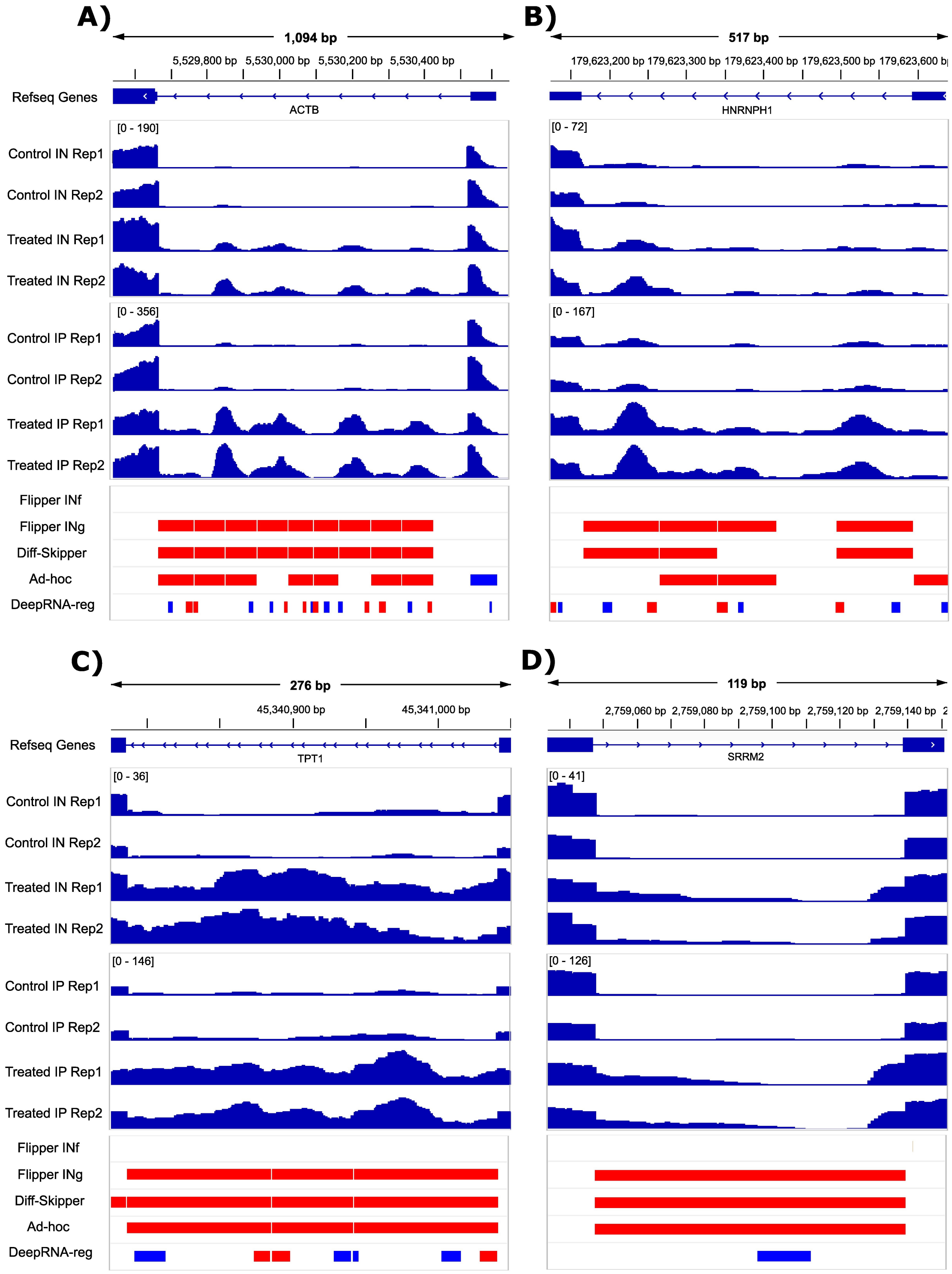


Fig. 4: Examples demonstrating the need for feature-level IN control at introns in (A) ACTB (also shown in Figure 3C), (B) HNRNPH1, (C) TPT1, and (D) SRRM2. In each case, local IN signal increases alongside IP signal. Detailed quantification of INf, INg, and intron retention is provided in Supplementary Tables 2 and 3.

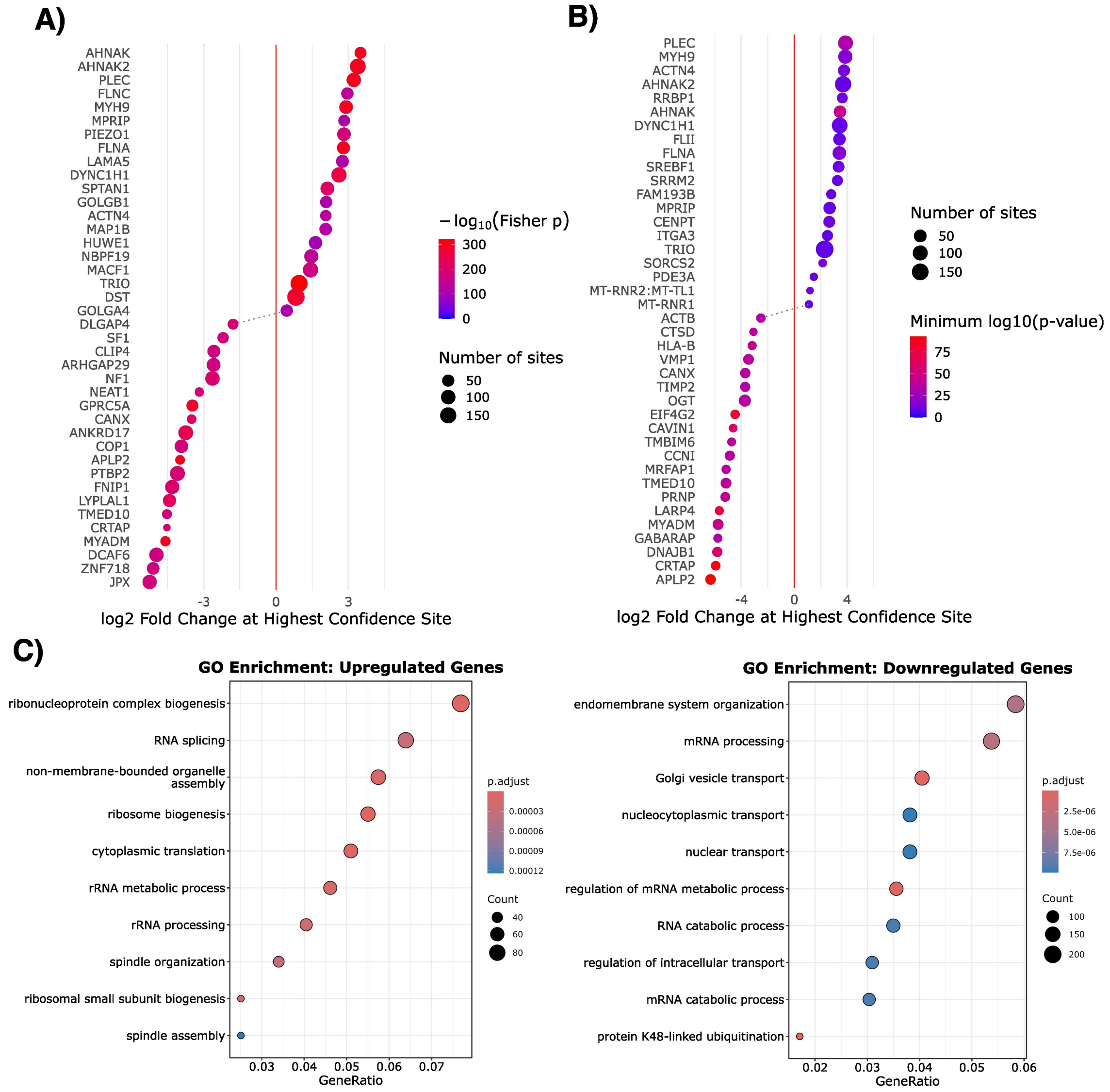


Fig. 5: Gene-level analyses performed by Flipper for the PUF60 eCLIP dataset. A) Dot plot showing the top 40 genes ranked by combined Fisher's p-value across all significant sites per gene, with total $\log_2$ fold change summed across sites. B) Dot plot showing the top 40 genes ranked by the minimum p-value among significant sites per gene, with the corresponding $\log_2$ fold change at that site. C) Gene Ontology (GO) analysis of genes containing significant binding sites, stratified by net positive (upregulated) or net negative (downregulated) fold change in binding.
